# Low support values and lack of reproducibility of molecular phylogenetic analysis of Copepoda orders

**DOI:** 10.1101/650507

**Authors:** Kirill V. Mikhailov, Viatcheslav N. Ivanenko

## Abstract

Reanalysis of the dataset used by Khodami *et al.* (2017) reveals low support values for the key nodes of the copepod (Crustacea) phylogeny and fails to reproduce the results obtained in the study. Maximum likelihood (ML) and Bayesian analyses with the dataset produce phylogenies that are inconsistent with the branching of copepod groups proposed by Khodami *et al.* (2017). The proposed phylogeny is refuted by the approximately unbiased (AU) statistical test, which undermines several conclusions drawn from the original study.

## Introduction

Copepoda is a large and diverse group of arthropods that encompasses roughly 10 well-recognized orders. To this day, the phylogenetic relationship between the copepod orders remains in dispute (Ferrari *et al.* 2010; Ferrari & von Vaupel Klein 2019; Ho 1994; Huys & Boxshall 1991; Khodami *et al.* 2019). Early molecular phylogenetic analyses of the copepod phylogeny based on rDNA sequences found it difficult to draw clear conclusions on the branching of several groups (Huys *et al.* 2007). In this context, the study by Khodami *et al.* (2017), which explored this question using additional phylogenetic marker genes (COI and histone H3) and a diverse sample of copepods, provides an important milestone for addressing the copepod phylogeny. The trees presented in the study featured high support values for almost all previously uncertain branches, thus portraying many of the open questions in copepod phylogeny as resolved. However, in our analyses using the dataset published by the authors, we have found it impossible to obtain high support values for many key branches of the tree or to simply replicate the inferred phylogeny. To confirm our negative results, we have conducted reanalysis of the original dataset made available by the authors using the methods indicated in the paper. Our analyses with both maximum likelihood and Bayesian approaches yield phylogenies that differ from the results presented in the study by Khodami *et al.* (2017). Furthermore, the phylogeny of copepods proposed by the authors is firmly rejected by the AU test.

## Results

The phylogenies reconstructed with the ML (Fig 1A and Supplementary Fig. S1) and Bayesian inference (Fig 1B and Supplementary Fig. S2) are in discord with the phylogenetic tree presented by Khodami *et al.* (2017) in fig. 2 (Fig 1C here). We do not observe the monophyly of Misophrioida in the reconstructed trees or the split of Harpacticoida into Polyarthra (Longipedidae and Canuellidae) and Oligoarthra. The Gelyelloida do not group sister to Harpacticoida and Cyclopoida. Postion of the divergent clade Monstrilloida is uncertain in the analyses – their sister group position to the Siphonostomatoida is only partially recovered in the Bayesian analysis. Generally, the phylogenetic trees obtained here have low support values, reflecting poor resolution of the relative branching of the copepod orders. This is in sharp contrast to high to maximal support values obtained in the ML and Bayesian analyses by Khodami *et al.* (2017 figs 2, 3, and S6).

**Fig. 1.**
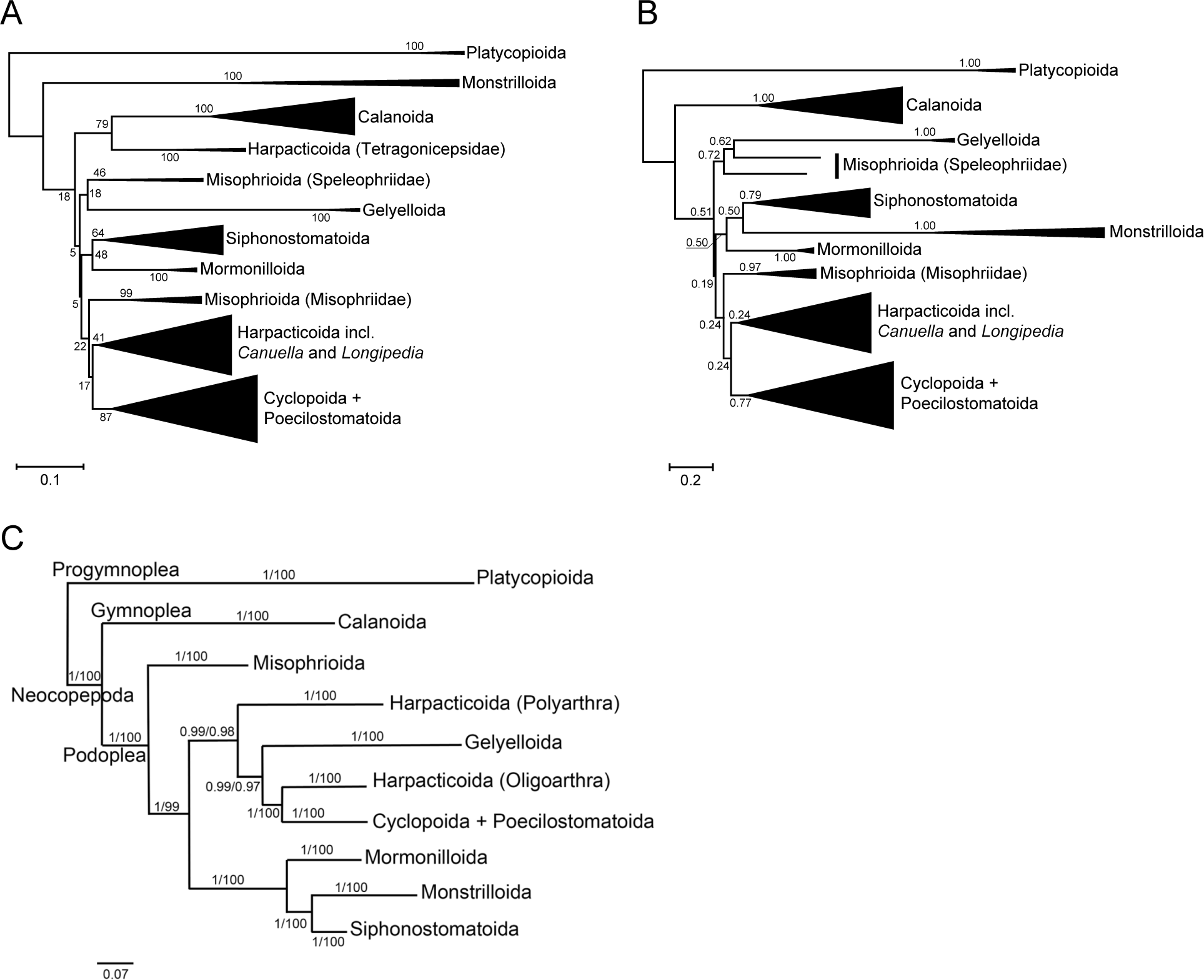
A. Phylogram of the copepod orders obtained in the RAxML analysis with the concatenated alignment of the four genes from the study by Khodami et al. (2017). Node support is indicated by the bootstrap values. B. Phylogram of the copepod orders recalculated with MrBayes using the concatenated alignment and summarized with a 25% burn-in after 20M generations. Node support is indicated by the posterior probabilities. C. Phylogram of the copepod orders from Khodami et al. (2017 fig 2): “Order-level phylogram of 10 copepod orders. Phylogenetic relationship of 210 copepod species (collapsed to order level) based on Maximum Likelihood and Bayesian analysis of 18S and 28S rRNA, COI mtDNA and H3 histone protein. Nodal support is indicated by posterior probabilities and bootstrap values.”

To confirm that the phylogenies obtained here and poor support values were not simply a result of insufficient tree reconstruction efforts, we conducted a test of alternative topologies using the trees obtained here and the tree from the original study by Khodami *et al*. (2017 fig. S6). The results of the AU test give preference to the ML tree topology over the alternatives (Table 1A). The *p*-value for the topology from the Bayesian analysis (Table 1B) is close to the 0.05 significance margin, scoring worse than the ML tree, presumably due to the differences in the evolutionary models used for tree inference. The tree topology from Khodami *et al.* (2017 fig. S6), however, scores significantly worse than the alternatives, and is rejected by the AU test at the 0.05 and 0.01 significance levels (Table 1C).

**Table 1.**
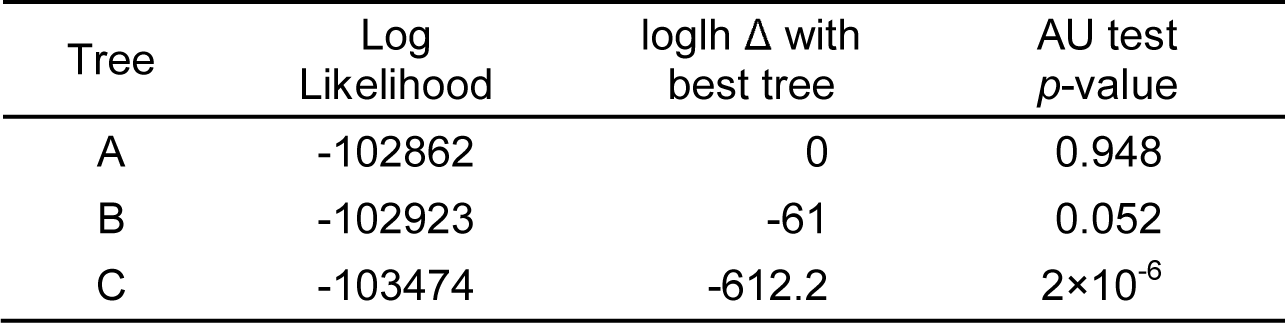
AU test of alternative tree topologies from Fig. 1: A – RaxML reconstruction, B – MrBayes reconstruction, C – tree topology from Khodami *et al.*, 2017 (figs 2 and S6).

| Tree | Log Likelihood | loglh $\Delta$ with best tree | AU test $p$-value |
| --- | --- | --- | --- |
| A | -102862 | 0 | 0.948 |
| B | -102923 | -61 | 0.052 |
| C | -103474 | -612.2 | $2 \times 10^{-6}$ |

## Discussion

Our findings demonstrate that the highly supported copepod phylogeny obtained by Khodami *et al*. (2017) is not reproducible. This indicates that the revision of copepod taxonomy undertaken by the authors of the original study is not supported by the molecular phylogenetic analyses. The rRNA genes (18S rRNA and 28S rRNA) are routinely used to propose phylogenetic relationships among copepod families or genera (Blanco-Bercial *et al.* 2011; Huys *et al.* 2006, 2012; Yeom *et al.* 2018), but these analyses fall short in resolving the relationship between copepod orders. The extended dataset does not significantly improve the resolution of copepod phylogeny, furthermore, the maximum likelihood and Bayesian analyses with the dataset fail to consistently support the monophyly of recognized orders Harpacticoida and Misophrioida. This lack of resolution might also be exacerbated by incomplete sampling of genes – the concatenated alignment produced by Khodami *et al.* (2017) does not equally cover all representatives of the copepod orders.

Low support values and a lack of reproducibility are not unusual problems. For example, Regier *et al.* (2010), cited approvingly by Khodami *et al.* (2017), attempted to analyze relationships among a large number of arthropod groups. Yet Rota-Stabelli *et al.* (2013), using data from Regier *et al.* (2010), were unable to resolve satisfactorily the relationships among pancrustaceans using nucleotide or amino acid sequences. They concluded that these relationships should be considered unresolved.

The work by Khodami *et al.* (2017) is an important contribution to the phylogenetic study of copepods; however, we advise that these results be treated with caution and emphasize that the issues with phylogeny of Copepoda are far from being settled.

## Methods

The phylogenetic analyses were performed with the concatenated alignment of four genes (18S rRNA, 28S rRNA, COI, and histone H3) between 203 copepod species, made available by the authors of the original study (Khodami *et al.* 2017) (https://treebase.org/treebase-web/search/study/matrices.html?id=20470, ID: M39845). Where possible, the tree reconstruction parameters were selected to parallel the settings used in the original study. The maximum likelihood analysis was carried out with RAxML using the GTRGAMMAI model optimized separately for the four gene partitions. The tree search in RAxML 8.2.9 (Stamatakis 2014) employed the random starting tree option (-d), and the tree node support values were evaluated with 10,000 bootstrap replicates. The Bayesian analysis was performed with MrBayes 3.2.6 (Ronquist *et al.* 2012). For rRNA gene partitions, we used the GTR model with 8 gamma-distributed rate categories and a proportion of invariants; for protein-coding gene partitions the codon model was used in conjunction with the GTR and 8 gamma-distributed rate categories. In the Bayesian analysis, the model parameters between the partitions were unlinked, and for the COI gene partition “metazoan mitochondrial” code was used. The analysis was performed in four independent runs of four chains each with 20M generations, and summarized with a 25% burn-in. For the alternative topology test, we used CONSEL (Shimodaira & Hasegawa 2001) with the site-wise likelihood values estimated by RAxML to compare the phylogenies obtained in this reanalysis with those of the original study. The site-wise likelihood values for the test were evaluated using the same model for RAxML as the one used for tree reconstruction. The CONSEL AU test was used for the estimation of *p*-values.

## Contributions

K.V.M. conducted data analysis and prepared manuscript; V.N.I. took part in planning, discussion of the results and preparing of manuscript.

