## Supplementary Fig. S1 and S2 for "Low support values and lack of reproducibility of molecular phylogenetic analysis of Copepoda orders"

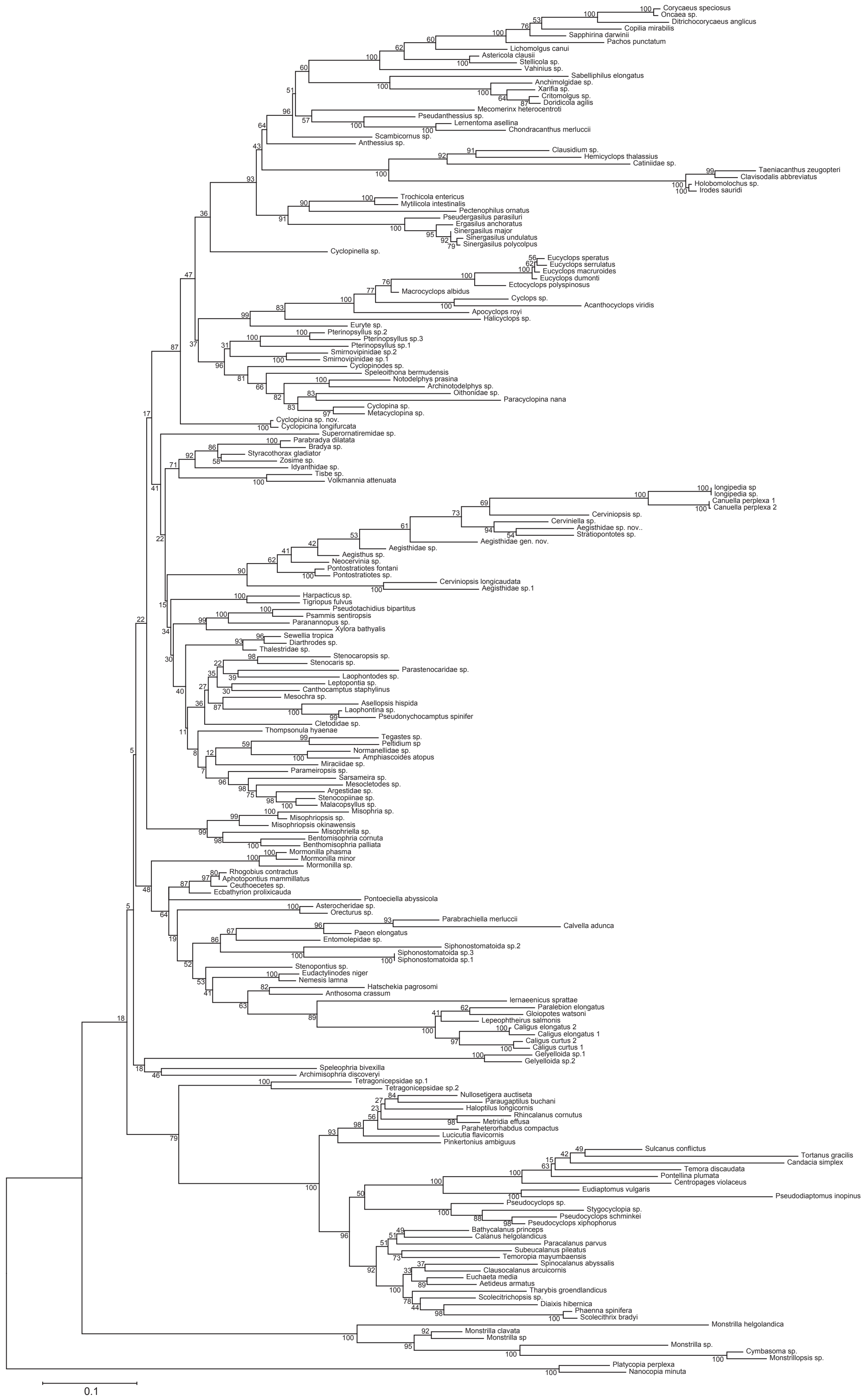


**Figure S1.** Maximum likelihood reconstruction with the concatenated alignment of the four genes (18S and 28S rRNA, COI mtDNA and H3 histone) from the study by Khodami et al., 2017. The tree was reconstructed with RAxML using the GTRGAMMAI model optimized separately for the four gene partitions; tree node support values are bootstrap percentages evaluated with 10,000 replicates.

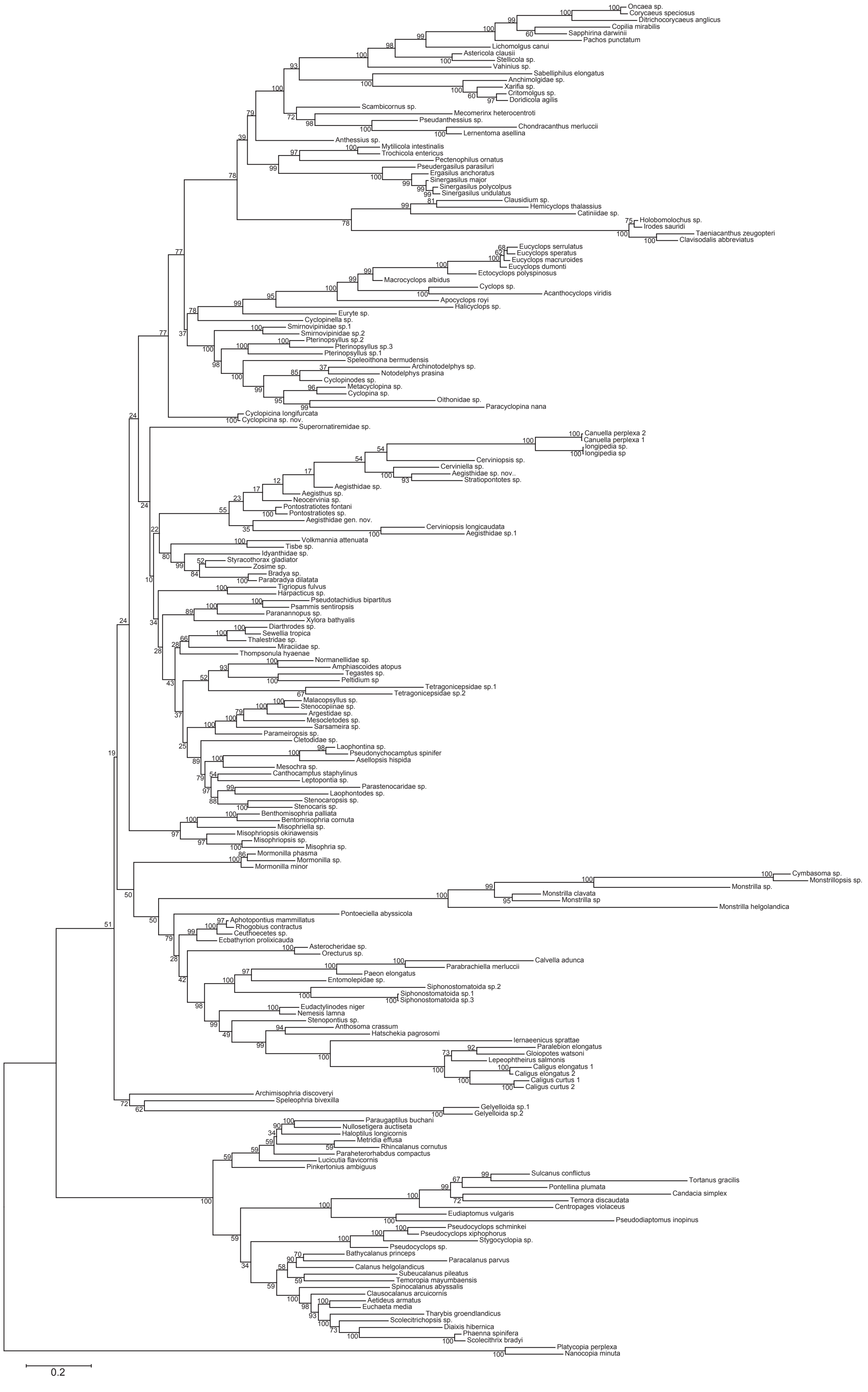


**Figure S2.** Bayesian inference with the concatenated alignment of the four genes (18S and 28S rRNA, COI mtDNA and H3 histone) from the study by Khodami et al., 2017. The tree was reconstructed by MrBayes using the GTR model for the rDNA partitions and the codon models in conjunction with GTR for the protein coding gene partitions; tree node support values are posterior probabilities given as percentage values.
